# Hazel dormouse in managed woodland select for young, dense, and species-rich tree stands

**DOI:** 10.1101/2022.04.06.487322

**Authors:** Rasmus Mohr Mortensen, Michelle Fyrstelin Fuller, Lars Dalby, Thomas Bjørneboe Berg, Peter Sunde

## Abstract

In fragmented forest landscapes, population persistence of arboreal species with limited dispersal ability may strongly depend on the quality of the remaining forest habitat. Using the hazel dormouse (*Muscardinus avellanarius*) as a model species, we studied habitat selection at two spatial scales (home range and within home range) in intensely managed woodlands at its northern distributional range in Denmark. We modelled selection at home range level as the conditional probability of occupancy of 588 nest boxes and nest tubes in 15 managed forests relative to habitat variables measured within 25 m radius. Habitat selection within home ranges was modelled by comparing habitat variables within 3 m radius of triangulated locations by 19 radio-tracked individuals (12 M, 7 F) when active at night with regularly distributed available locations within their home ranges.

At both spatial scales, hazel dormice strongly selected sites with high abundance-weighted species richness and high vegetation density of woody plants. On home range level, they furthermore selected for young tree vegetation, while they within home ranges selected for intermediate aged tree stands (maximum trunk circumference: 1.50 m). The predicted probability of presence in nest boxes or nest tubes varied from less than 1% to more than 99% as a combined function of three habitat variables. From May to October, selection for abundance-weighted species richness of woody plants of radio-tagged individuals decreased with date and body weight, suggesting that a diverse food base is particularly important early in their season of activity and for lean and small (growing) individuals. Selection for dense vegetation increased with body mass and mean available vegetation density within home ranges, indicating behavioural variability related to changes in energy expenditure and need for safety among individuals.

The study demonstrates that the hazel dormouse has specific habitat requirements related to food and safety that can be accommodated with relatively simple means in managed forests.

**Graphical abstract:** 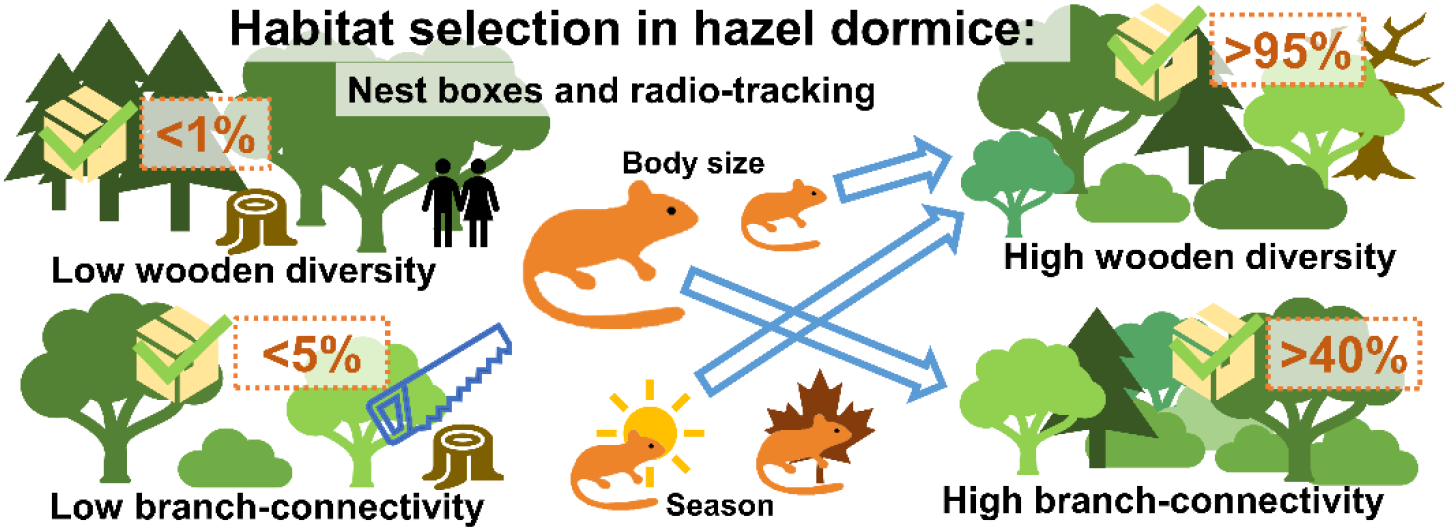

## 1. Introduction

Large parts of terrestrial ecosystems are heavily influenced by human activities including forestry practices (Bengtsson *et al*., 2000; Williams *et al*., 2020), which through loss, modification, and fragmentation of natural forest habitats have led to substantial changes in forest structure and dynamics (Paillet *et al*., 2010; Burrascano *et al*., 2013). Historically, various forestry practices have caused a simplification of forest ecosystems to promote production (Nilsson, 1997; Kaplan *et al*., 2009; McGrath *et al*., 2015), consequently affecting several sensitive and narrow-range species that depend on structures and processes of old-growth forests (Paillet *et al*., 2010). Therefore, most of the current forests in Europe lack natural variation and ecological continuity giving little room for natural open structures and trees of various succession state (Nilsson, 1997; Bengtsson *et al*., 2000; Nordén *et al*., 2014).

Anthropogenic activities in the forest alter the distribution and abundance of resources, predators, and social interactions (Bengtsson *et al*., 2000), which consequently affect the movements and habitat use of wild forest-dwelling animals as habitats may become unavailable or less favourable (Gallagher *et al*., 2017). Forest habitats are home to many protected and threatened animal species that may be adversely affected by various forest management activities (Danneyrolles *et al*., 2019), which potentially can have both short-term effects on individuals of various life-stages and with various life history strategies, as well as long-term effects on population dynamics (Blumstein, 2010; Mortensen and Rosell, 2020). This makes it a legal imperative for private and public forest owners to reduce or mitigate potential harm caused by forestry activities (Young *et al*., 2005). To counteract loss of biodiversity and protect endangered forest-dwelling animals that depend on specific forest conditions for e.g. foraging and breeding, active management is often needed (Bauhus *et al*., 2009), and forest management have to take these ecologically important forest habitats into account. However, the responses of protected species to various forestry activities are often not well understood and critical habitats may not be well-known (Nordén *et al*., 2014). Consequently, the effects of many forestry activities may often be assumed with a general approach of applied measures. This indicates the importance of strong links between research and practice to improve quality and validity of conservation management plans (Lindenmayer, 1999; Bergès and Dupouey, 2021). Although it is challenging to quantify which habitats animals in an intensely managed forest landscape have available to them and to what extent they make use of them, this information is needed to determine the habitat value for the animal and put confounding ecological variables affecting this into context (Bleicher and Rosenzweig, 2018). Furthermore, this knowledge will prove helpful when designing management plans and employing resources most efficiently (Petersen et al., 2016).

In this study, we examine habitat selection at two spatial scales in an arboreal mammal inhabiting intensely managed woodland using the hazel dormouse (*Muscardinus avellanarius, dormice hereafter*) as a model species. The hazel dormouse, is a small (adults typically weigh 10-35 g), nocturnal arboreal rodent with a geographic range covering large parts of Europe from the Mediterranean in the south to the southern parts of Scandinavia in the North (Juškaitis, 2014a). Although widespread, the hazel dormouse is considered a threatened species in large parts of its distribution (Vilhelmsen, 2003; Temple and Terry, 2007; Juškaitis, 2014a) due to its sensitivity to habitat loss, habitat fragmentation, and unfavourable forest management practices (Mortelliti *et al*., 2014; Ramakers *et al*., 2014; Sozio *et al*., 2016). However, despite receiving attention and several conservation measures, populations keep declining in some areas, emphasizing the need to improve our understanding of what drives the ecological dynamics of hazel dormouse populations (Goodwin *et al*., 2017; Fedyń *et al*., 2021).

Hazel dormice are typically associated with dynamic forest habitats with high plant diversity, trees of various ages, and enough light allowing a rich understory and regeneration to take place (Bright and Morris 1996). These conditions seem to favour the dormouse by providing resting and breeding places, as well as vegetation for foraging and movement. The dormouse is dependent on a continuous food supply of flowers, fruits, fungi, and invertebrates from the beginning of its active period in the spring until it hibernates in the winter (Bright and Morris, 1996; Juškaitis and Baltrūnaitė, 2013; Juškaitis *et al*., 2016; Büchner *et al*., 2018; Goodwin *et al*., 2020). Favoured vegetation types have shown to vary between different habitats, suggesting that dormice are quite adaptable in their selection for food items and may choose different trophic levels depending on seasonal phenological change (Juškaitis, 2007; Juškaitis and Baltrūnaitė, 2013; Chanin *et al*., 2015; Goodwin *et al*., 2020). A well-structured dense vegetation with high branch-connectivity between trees and shrubs enables safe movement options in protection from predators (Bright, 1998; Juškaitis *et al*., 2013) as dormice generally seem to avoid crossing open ground, although studies have shown that long distance field crossings can occur (Büchner, 2008; Mortelliti *et al*., 2013). The need for dynamic and successional wooded habitats exemplifies the challenges when wanting to conserve a species in a system that is subject to frequent management and alternation of habitats as the dormice are likely to require active management to maintain their favoured habitats and facilitate persistence of dormouse populations. Studies have shown that active management of woodland habitats increases survival and body condition and dormouse populations have shown to be more resilient (Trout *et al*., 2012; Juškaitis, 2014a; Sozio *et al*., 2016; Goodwin *et al*., 2018b).

As a strictly arboreal rodent with low recruitment rate and low population densities (Bright and Morris, 1996; Büchner *et al*., 2003; Juškaitis, 2014a, b) the dormouse is particularly vulnerable to habitat fragmentation and habitat loss that follows intense management of woodlands (Trout *et al*., 2012). In regions where forest is sparse, fragmented, and managed, silvicultural management practices may be of great importance for the density and ultimately viability of the remaining dormouse populations (Mortelliti *et al*., 2011; Zapponi *et al*., 2013; Mortelliti *et al*., 2014; Dondina *et al*., 2016). There is a need for improved evidence-based knowledge on the habitat requirements of the hazel dormouse in heavily managed woodlands to improve conservation and management options of potential dormouse forest habitats (Cartledge *et al*., 2021). From a legal perspective, knowledge of optimal habitat features for dormice is of particular importance for national management authorities that are obliged to protect the species throughout the EU where the species is placed in the Annex IV of the EU Habitats Directive 92/43/EEC. In particular, this applies to a country like Denmark, where the species is declining and exists on the northern limit of its geographical range in five isolated populations, some of which seem to extend to a few square kilometres of managed forest where the species is reported to be rare and hard to locate (Therkildsen *et al*., 2020). As several of these populations may be too small to be viable in the long term, increasing the ecological capacity through habitat improvements may be crucial to prevent these populations becoming extinct.

We studied the habitat selection of hazel dormice in its remaining population strongholds in Denmark (all intensively managed woodlands) at two spatial scales. On the first scale, we assessed the conditional probability of presence (home range level) within known population areas in terms of probability of occupancy of nest boxes and nest tubes relative to habitat variables measured within a 25 m radius. On the second scale (within home range) we compared habitat variables of locations used by radio-tagged individuals with habitat variables of regularly distributed available locations within the home ranges. We hypothesized that on both spatial scales, dormice would select for habitat features associated with a rich and diverse food base (species abundance score of soft mast species, hard mast species, coniferous species, or other woody species of interest), and safety and climbing ability (tree vegetation density).

Furthermore, we investigated how habitat selection varied between individuals (sex, body size, time of the year, and mean habitat composition within home range).

## 2. Materials and methods

### 2.1 Occupancy of home ranges within populations

To quantify occupancy of potential hazel dormouse home ranges within populations in relation to habitat composition, we registered the frequency by which nest boxes and nest tubes were used by dormice at various forest locations across Denmark where dormice occurred. Nest boxes have been put up in several Danish forests to improve the conditions for the hazel dormouse (Vilhelmsen, 2003) and are typically examined yearly for nests. As part of a national monitoring program (Søgaard *et al*., 2015; Kjær *et al*., 2021), nest tubes were furthermore placed throughout the country in April-November 2012 and April-December 2013, and examined for presence of dormice when taken down before the winter season.

Our study included 588 nest boxes and nest tubes from fifteen locations from the two largest Danish population areas (Fig. 1). Dormice were known to be present at the locations, as evident from presence of at least one nest box/tube with clear signs of hazel dormouse presence (Morris *et al*., 1990). The response variable was as of whether the nest box/tube had been used or not used by dormice by the end of the census year. All nest boxes and nest tubes were placed in managed forest habitats presumed to be optimal for dormice after Danish habitat standards.

**Figure 1.**
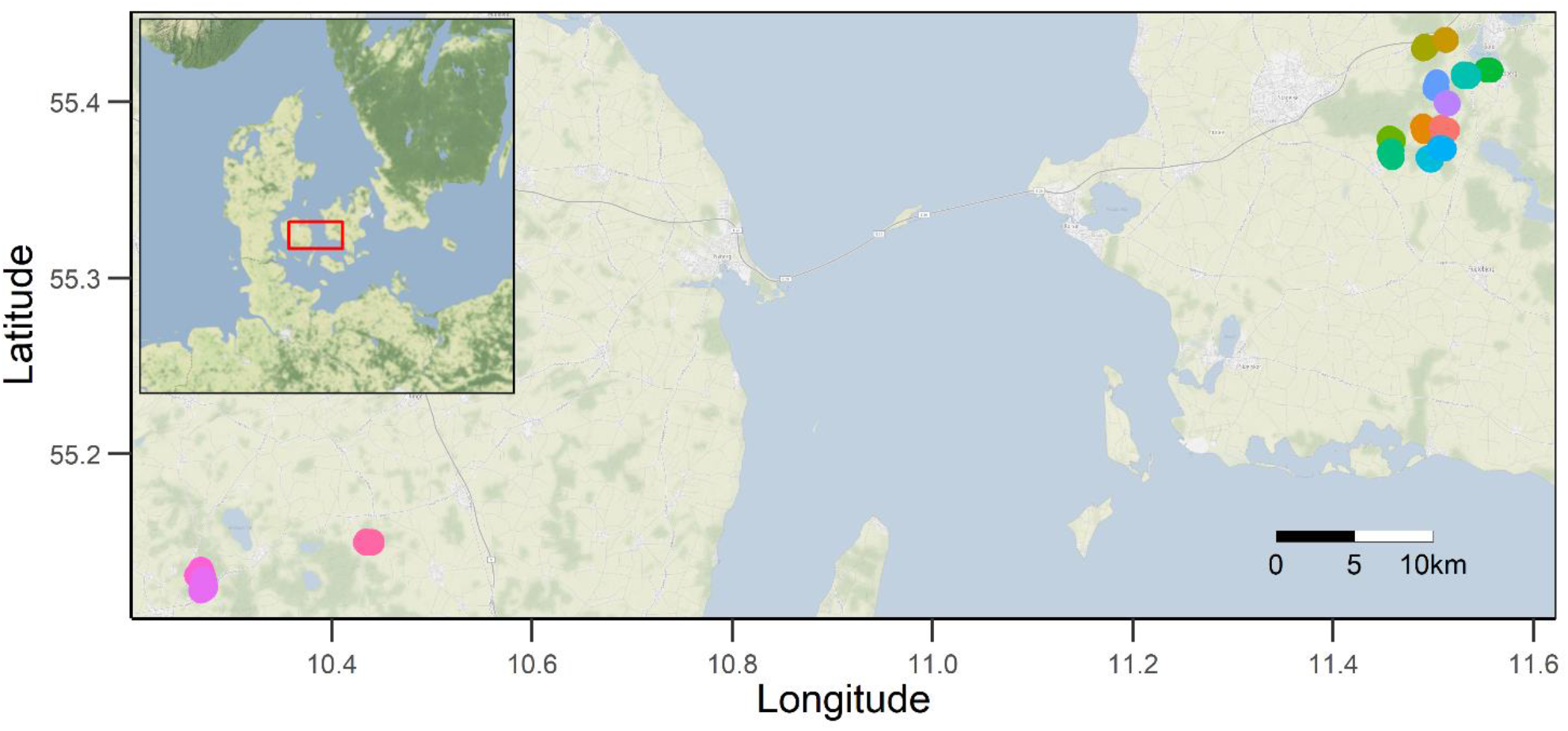
Overview of the 15 managed forest locations which were surveyed for hazel dormouse presence using nest boxes and nest tubes in 2012-2013. Red box on the map of Denmark shows the location of the zoomed map view.

### 2.2 Telemetry

To study how hazel dormice use habitats within their home ranges, we radio-tagged 20 individuals (of which telemetry data was obtained from 19) from a population in a managed forest located in Svanninge Bjerge, Denmark, (55º07’N 10º16’E) from May to October 2013 (4 F, 9 M) and from June to July 2014 (4 F, 3 M) (Table S1). Individuals were caught in nest boxes at daytime. They were sexed and weighed after which a 0.39-0.43 g VHF transmitter (PIP3 Ag317, Biotrack Ltd.) was glued onto a shaved patch on the back of the individual. The tags dropped off the animal after 1-8 days and their weights never exceeded 4% of the body mass. Capture and handling were done as swiftly and gently as possible to reduce short- and long-term effects (Mortensen and Rosell, 2020). No captured individuals were injured during capture and handling, and they were all released back into their nest box after approximately 20 minutes of handling. Capture, handling, and tagging were licensed through a general institutional permission (Aarhus University, Department of Ecoscience) for capturing and marking birds and mammals, issued by the Danish Nature and Forest Agency (reference: SM 302-009) and our study met the ASAB/ABS Guidelines for the treatment of animals in behavioural research and teaching (Buchanan *et al*., 2012).

The dormice were radio-tracked continuously from before sunset to after sunrise by means of triangulation on approximately 5-50 m distance from fixed bearing points in the terrain. Temperature, precipitation, wind, and light intensity were noted simultaneously. Each triangulation attempt was performed approximately 10 to 20 minutes apart, but with longer breaks when individuals made swift moves from one location to another. Using the triangulation fixes, we calculated 95% autocorrelated kernel density estimates (AKDE) to estimate the area of activity (home range) of each individual (Fleming *et al*., 2015). Regularly distributed locations in a 10-m grid within the home range of each individual were used to quantify the available habitat distribution for each individual (Fig. 2).

**Figure 2.**
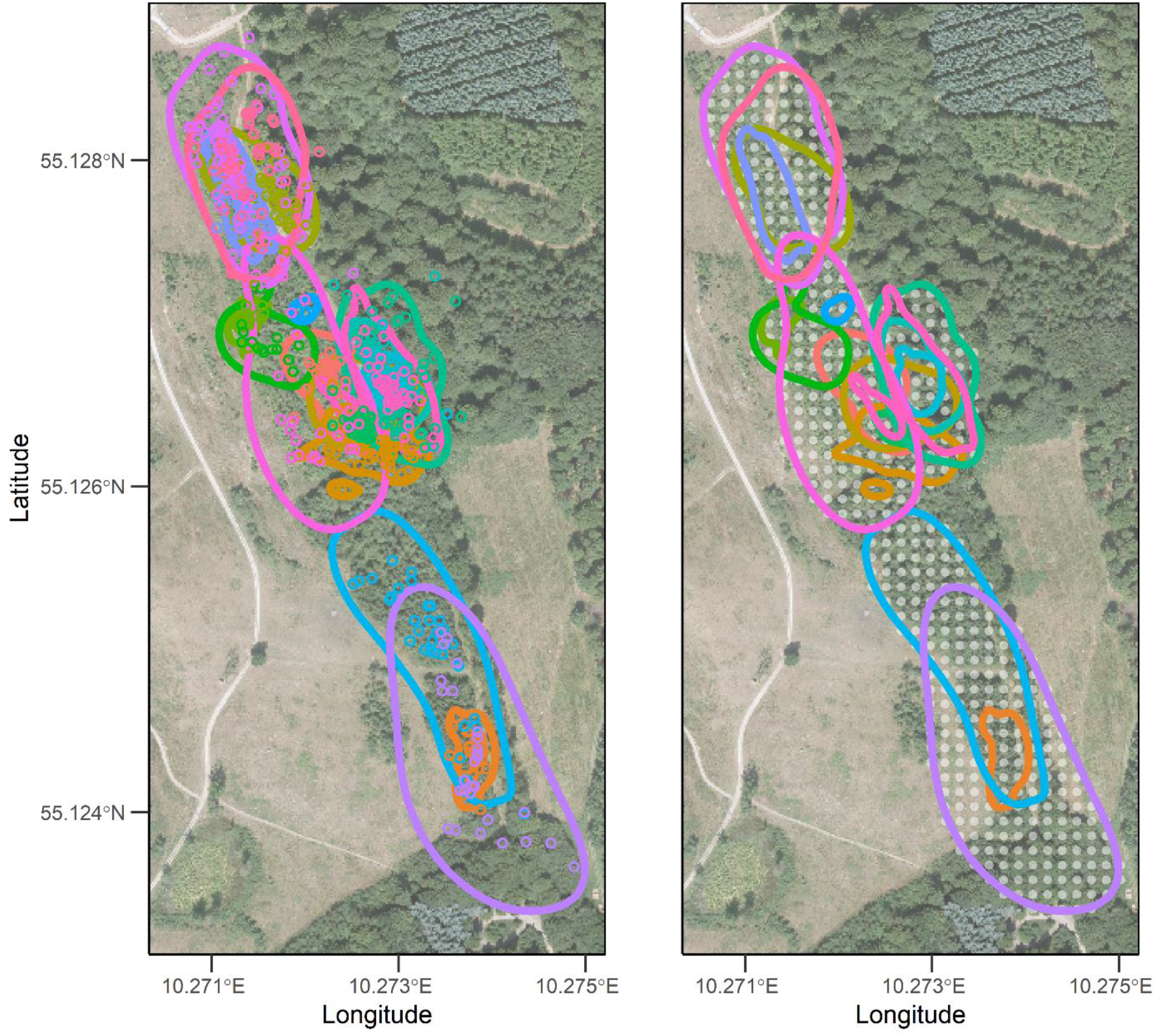
Nocturnal telemetry locations (coloured points) and derived home ranges (95% autocorrelated kernel density isopleths) of 19 radio-tagged dormice in Svanninge Bjerge, Denmark, 2013-2014. Grey dots indicate regularly distributed (10-m distance) available location (grey points) within each dormouse’s home range.

### 2.3 Assessment of habitat variables

From August to October 2013, we visited all nest boxes and tubes (or their coordinates if taken down the previous year) and registered habitat characteristics of presumed importance for dormice (Table 1) within a 25 m radius (the approximate extension of a dormouse home range in Denmark: Fig. 2). To quantify the habitat selection within home ranges, we revisited each triangulated telemetry location at daytime in the days following radio-tracking and assessed habitat variables (Table 1) within a 3 m radius. To assess the distribution of available habitats, we also assessed regularly distributed locations in a 10-m grid within the home range of each individual (Fig. 2).

**Table 1.** Description of habitat variables assessed within a 25 m radius from nest boxes or nest tubes (analysis of location of home ranges within populations, second order selection) and a 3 m radius from triangulated locations and regularly distributed “availability locations” within the home ranges of tracked hazel dormice (analysis of habitat selection within home ranges, third order selection). Variable name in parentheses.

| Variable | Definition |
| --- | --- |
| Tree canopy cover (Canopy) | Percentage of tree canopy cover within either 25 m or 3 m. |
| Tree height (Height) | Height (m) of highest tree in the assessment area. |
| Tree girth (Girth) | Circumference (m) of the thickest tree trunk in the assessment area. |
| Vegetation density (VD) | In three different vertical layers (Low: 0-2 m, Middle: 2-10 m, High: >10 m), the densest sight line from the centre to the edge of the assessment circle were scored as 1: open vegetation (gaps >2 m), 2: spread vegetation (gaps 1-2 m), 3: moderately dense vegetation (gaps <1 m), 4: dense vegetation (no gaps). In the analyses we included vegetation density scores for each vertical layer ($VD_{Low}$, $VD_{Middle}$, $VD_{High}$) and averaged combinations over the layers ($VD_{LM}$, $VD_{MH}$, $VD_{All}$). |
| Abundance score (AS) | Every species of woody plants detected in the assessment area were given an abundance score: 0: species is absent, 1: species is present, 2: species is abundant (readily observable) in >25 % of the assessment area, 3: species is (partly) dominant in >25% of the area within an assessment radius of 25 m |
|  | (second order analysis) or >50 % of the area within an assessment radius of 3 m (third order analysis). |
| Summed species abundance score (SSAS) | The abundance score summed over all species of woody plants (SSAS). The index was calculated for all species of woody plants (SSAS<sub>all</sub>), all soft mast species of woody plants (SSAS<sub>softmast</sub>), all hard mast species of woody plants (SSAS<sub>hardmast</sub>), all coniferous species (SSAS<sub>conifer</sub>), and for woody species that were selected by the hazel dormice (SSAS<sub>selected</sub>). |
| Soft mast species: | Species that produce berries and fleshy fruits (aggregates, pomes, and drupes):<br>Apple (*Malus sylvestris*), blackberry (*Rubus plicatus*), buckthorn (*Frangula alnus*), cherry (*Prunus* spp.), dogwood (*Cornus* spp.), elder (*Sambucus* spp.), guelder rose (*Viburnum opulus*), hawthorn (*Crataegus* spp.), holly (*Ilex aquifolium*), honeysuckle (*Lonicera* spp.), horse chestnut (*Aesculus hippocastanum*), linden (*Tilia cordata*), raspberry (*Rubus ideaus*), rowan (*Sorbus* spp.), and sloe (*Prunus spinosa*). |
| Hard mast species | Species that produce nuts and winged seeds:<br>Alder (*Alnus* spp.), ash (*Fraxinus excelsior*), beech (*Fagus sylvatica*), birch (*Betula* spp.), elm (*Ulmus* spp.), hazel (*Corylus avellana*), maple (*Acer* spp.), and oak (*Quercus* spp). |
| Coniferous species | Species that produce strobili:<br>Fir (*Abies* spp.), larch (*Larix* spp.), pine (*Pinus* spp. and *Pseudotsuga menziesii*), spruce (*Picea* spp.), and thuja (*Thuja plicata*). |
| Capsules and legumes | Species that produce capsules and legumes: |
|  | Aspen (*Populus* spp.), broom (*Cytisus scoparius*), dog rose (*Rosa canina*), spindle (*Euonymus europaeus*), and willow (*Salix* spp.). |

At each location, we recorded all species of woody plants (trees above 1.5 m and bushes above 0.5 m) and their abundance in the assessment area were estimated on a species abundance index from 0 to 3 (abundance score, AS: Table 1). As a weighted index for species richness and abundance of woody species, we used summed species abundance score (SSAS: Table 1) of the various woody species. This index correlated highly with species richness (r > 0.9), but furthermore integrated the abundance of the species registered. In addition to the overall SSAS based on all woody species, we also calculated SSAS-scores of species groups that may be particularly preferred or avoided by the dormice: hard mast, soft mast, coniferous, capsules and legumes (Table 1).

For three different vertical layers (Low: 0-2 m, Middle: 2-10 m, High >10 m) the horizontal vegetation density was scored as the shortest sight line from the centre to the edge of the assessment circle from which a person would be decently visible (index ranging from 1 [open] to 4 [dense], Table 1). Furthermore, light incidence was estimated as percentage tree canopy cover of the assessment circle. As a proxy of forest age and succession stage of the various sites, we chose to estimate the tree height of the tallest tree and use a measuring tape to measure the circumference at breast height of the thickest tree trunk within the assessment area.

### 2.4 Statistical analysis

We investigated occupancy within populations (second order selection, Johnson (1980)) by comparing habitat variables (Table 1) of used and unused nest boxes and nest tubes in fifteen managed forest locations in Denmark (Fig. 1). The conditional probability of a nest box or tube being occupied was modelled in a resource selection probability function (RSPF) (Lele *et al*., 2013) using logistic regression with logit link and census location as a random intercept. As every resource unit (nest box or nest tube) could be categorised as either used (present) or unused (absent), the RSPFs not only estimated selection but also the absolute conditional probability that a nest box or nest tube would be occupied as function of habitat composition.

To reduce the number of covariates in the RSPF and avoid overfitting (Jenkins and Quintana-Ascencio, 2020), we tested the effect of abundance (score from 0 [not present] to 3 [dominant]: Table 1) of different woody plant species in separate models for each species registered in a minimum of 30 of the 588 locations. As an overall measure of the composite abundance of all woody species, we furthermore included the summed species abundance score (SSAS) of these selected woody plant species in the predictive model. As structural variables, we also tested indices for vegetation density (horizontal sight in three different layers: Table 1) and maturity (circumference of the thickest trunk at breast height: Table 1)

We investigated the habitat selection within home ranges (third order selection, Johnson (1980)) by comparing habitat variables (Table 1) of triangulated telemetry locations from night (when the hazel dormice were active) with habitat variables of regularly distributed available locations within the home range of each individual (Fig. 2). The relative probability of an area with given habitat variables being selected compared to areas the dormouse had available during the tracking period was modelled in a resource selection function (RSF) (Avgar *et al*., 2017; Northrup *et al*., 2022) using logistic regression with logit link and individuals as a random effect. As in the second order analysis, we analysed what woody plant species the hazel dormice selected in a separate model and included the SSAS of the selected woody plant species in the RSF together with the other covariate groups. In addition, we examined the context dependent selection in individual-specific models weighed by the inverse-variance of the coefficient estimates by fitting the RSF with no random effect to each individual (Gillies *et al*., 2006; Muff *et al*., 2020). This enabled us to investigate how ecological variations between individuals (sex, body size), season (time of the year), and home range compositions (home range size, mean vegetation density, mean tree canopy cover, mean height of highest tree, mean circumference of thickest tree, and mean SSAS) may affect the found resource selection functions (Mysterud and Ims, 1998; Gillies *et al*., 2006; McLoughlin *et al*., 2010; Ariano-Sánchez *et al*., 2020; Mortensen *et al*., 2021).

In all analyses, a priori lists of candidate models were defined based on ecologically relevant combinations of fixed effects to account for variability in endogenous (such as sex and body mass) and exogenous (such as home range size and composition) that may be important in describing the ecology of the hazel dormouse. The fixed effects used in all analyses were not strongly correlated (Pearson r coefficients < 0.6) and variance inflation factor values were less than 3 (Zuur *et al*., 2009). Model selection was based on Akaike’s information criterion corrected for small sample size (AICc) (Burnham and Anderson, 2002) and carried out using the R packages ‘glmmTMB’ v. 1.0.2.1 (Magnusson *et al*., 2017) and ‘MuMIn’ v. 1.43.17 (Barton, 2018). The most parsimonious models within ΔAICc < 2 were chosen as the best models to describe the variation (Burnham and Anderson, 2002; Arnold, 2010). Lists of top candidate models for all analyses can be found in the supplemental material. The best models were visually validated using the R package ‘DHARMa’ v. 0.4.1 (Hartig, 2017) to plot standardised model residuals against the fitted values (Zuur *et al*., 2009). In the most parsimonious models, variables that included zero within their 95% confidence interval (CI) were reported as unclear effects (Arnold, 2010; Muff *et al*., 2021). Confidence intervals were bootstrapped with 10,000 simulations to obtain robust estimates (Fieberg *et al*., 2020). All analyses were conducted in R 4.1.0 (R Core Team, 2021).

## 3. Results

### 3.1 Location of home ranges within populations

We found signs of hazel dormouse occupancy in 191 of 588 nest boxes. The most parsimonious nest box occupancy model included habitat variables for vegetation density, SSAS, and vegetation age (Table 2, Table S2). Within the observation range, the predicted probability of presence in nest boxes or nest tubes varied from less than 1% to more than 99% as a combined function of the three habitat variables: vegetation density score below 10 m (positive, Fig 3a), SSAS of selected woody species (positive, Fig. 3b), and circumference of thickest trunk (negative, Fig. 3c). The most powerful single predictor was SSAS of selected woody species that predicted a variation of probability of occupancy from less than 1% to more than 95% (Fig. 3b).

**Table 2.**
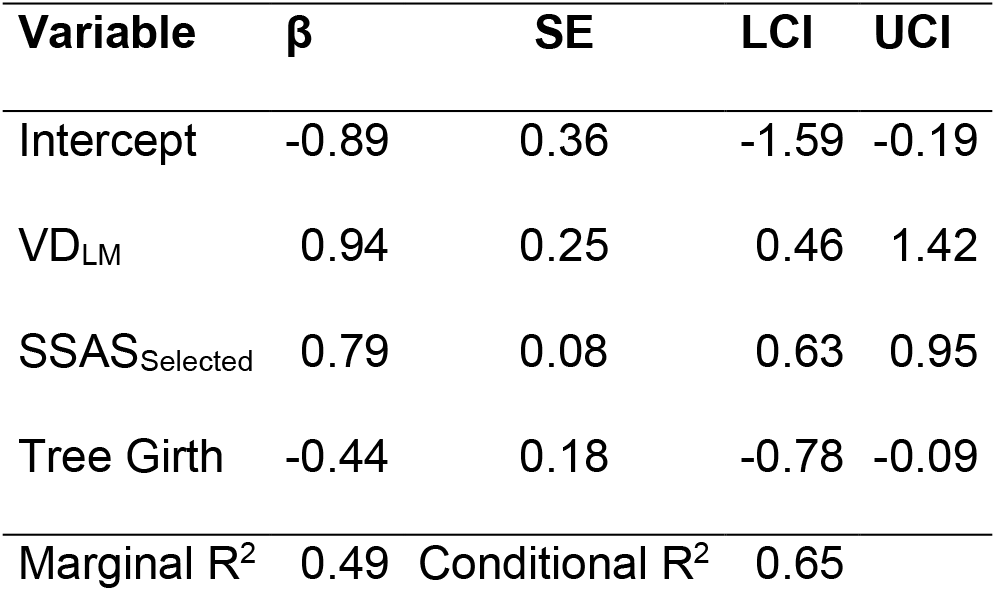
Effect size (β), standard error (SE), lower (LCI) and upper (UCI) 95% confidence interval of explanatory variables for the analysis of hazel dormouse occupancy in nest boxes and nest tubes (n = 588) across dormouse populations in fifteen managed forest patches in Denmark.

| Variable | $\beta$ | SE | LCI | UCI |
| --- | --- | --- | --- | --- |
| Intercept | -0.89 | 0.36 | -1.59 | -0.19 |
| VD<sub>LM</sub> | 0.94 | 0.25 | 0.46 | 1.42 |
| SSAS<sub>Selected</sub> | 0.79 | 0.08 | 0.63 | 0.95 |
| Tree Girth | -0.44 | 0.18 | -0.78 | -0.09 |
| Marginal R<sup>2</sup> | 0.49 | Conditional R<sup>2</sup> | 0.65 |  |

**Figure 3.**
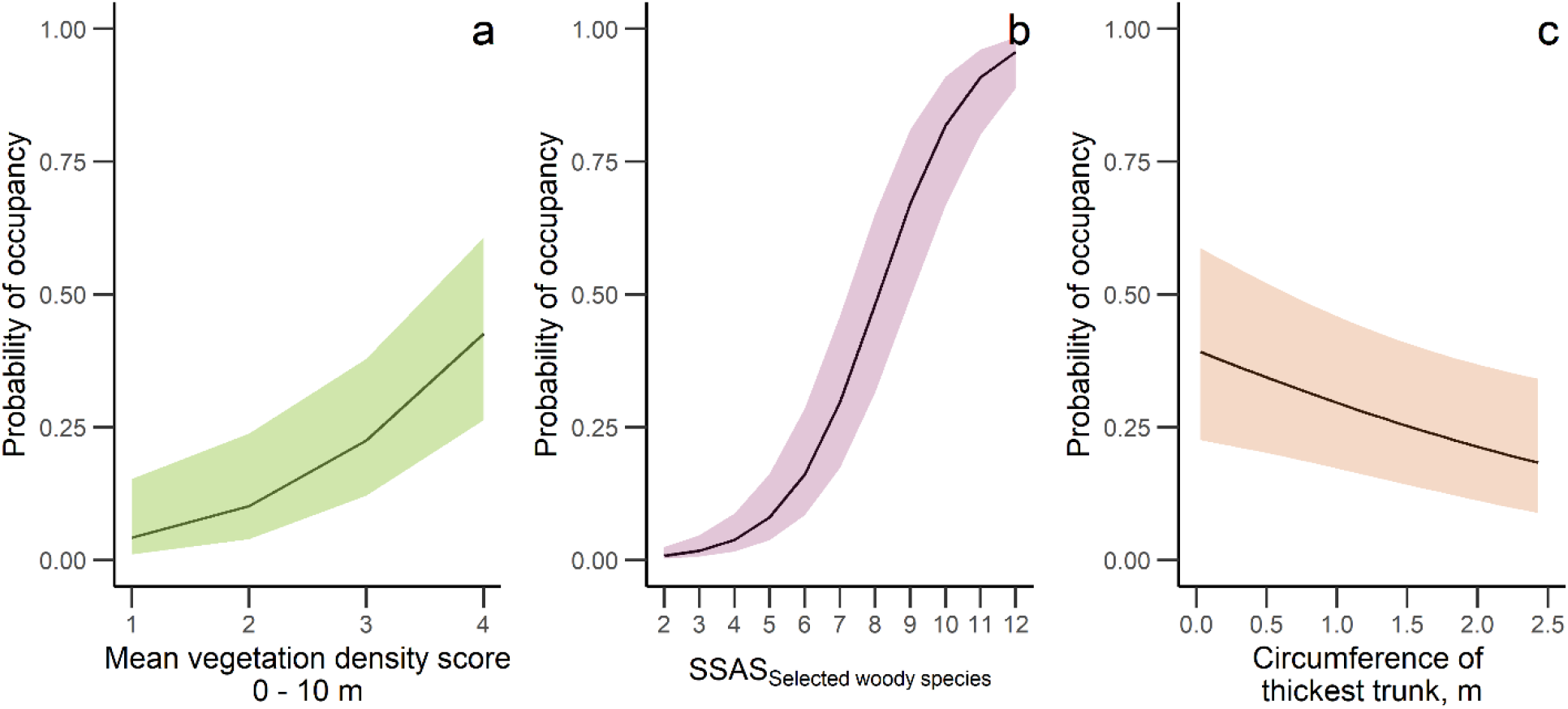
The predicted relationship with 95% confidence zones between probability of hazel dormouse occupancy in nest boxes and nest tubes at 15 managed forest patches in Denmark and (a) mean vegetation density score below 10 m, (b) summed species abundance score (SSAS) of woody species selected by the dormice, and (c) circumference of the thickest trunk.

Nest box and nest tube use was highly conditional on the presence of specific woody species groups as it increased significantly with summed abundance of blackberry (*Rubus plicatus*), beech (*Fagus sylvatica*), pine (*Pinus* spp. and *Pseudotsuga menziesii*), hazel (*Corylus avellana*), elder (*Sambucus* spp.), larch (*Larix* spp.), willow (*Salix* spp.), rowan (*Sorbus* spp.), and hawthorn (*Crataegus* spp.) (Table 3, Table S3, Fig. 3b). These woody species appeared more important for the hazel dormouse’s nest box selection than hard mast species or soft mast species alone, or than overall summed abundance of woody species near the nest box or nest tube (Table S2).

**Table 3.** Effect size (β), standard error (SE), lower (LCI) and upper (UCI) 95% confidence interval of explanatory variables for the analysis of hazel dormouse occupancy in nest boxes and nest tubes (n = 588) across dormouse populations in fifteen managed forest patches in Denmark as a combined function of abundance scores of ten woody species within a 25 m radius of the assessment nest boxes and nest tubes.

| Variable | $\beta$ | SE | LCI | UCI |
| --- | --- | --- | --- | --- |
| Intercept | -0.84 | 0.37 | -1.60 | -0.11 |
| Blackberry (*Rubus plicatus*) | 0.87 | 0.20 | 0.49 | 1.26 |
| Beech (*Fagus sylvatica*) | 0.80 | 0.17 | 0.46 | 1.14 |
| Pine (*Pinus* spp. and *Pseudotsuga menziesii*) | 1.11 | 0.40 | 0.32 | 1.89 |
| Spruce (*Picea* spp.) | 0.29 | 0.15 | 0.00 | 0.58 |
| Hazel (*Corylus avellana*) | 0.40 | 0.17 | 0.08 | 0.73 |
| Elder (*Sambucus* spp.) | 0.80 | 0.33 | 0.16 | 1.45 |
| Larch (*Larix* spp.) | 0.50 | 0.17 | 0.16 | 0.85 |
| Willow (*Salix* spp.) | 1.00 | 0.19 | 0.62 | 1.38 |
| Rowan (*Sorbus* spp.) | 1.23 | 0.20 | 0.85 | 1.61 |
| Hawthorne (*Crataegus* spp.) | 1.06 | 0.28 | 0.52 | 1.61 |
| Marginal R<sup>2</sup> | 0.47 | Conditional R<sup>2</sup> | 0.64 |  |

### 3.2 Habitat selection within home ranges

Nineteen hazel dormice were radio-tracked for a total of 73 tracking nights (1 to 8 nights per individual, mean ± SD = 4.1 ± 2.0 nights), resulting in a total of 953 telemetry fixes (50 ± 29 per individual, Table S1). The most parsimonious model included habitat variables for vegetation density (positive: Fig. 4a), SSAS of selected woody species (positive; Fig 4b), and vegetation age (humpbacked selection with the highest selection for tree stands with circumferences of the thickest trunk ∼ 1.5 m: Fig. 4c) (Table 4). Of individual species of woody plants, hazel dormice selected for maple (*Acer pseudoplatanus*), honeysuckle (*Lonicera periclymenum*), elder (*Sambucus* spp.), rowan (*Sorbus* spp.), hazel (*Corylus avellana*), birch (*Betula* spp.), beech (*Fagus sylvatica*), raspberry (*Rubus idaeus*), cherry (*Prunus avium*), willow (*Salix* spp.), and blackberry (*Rubus plicatus*) (Table 5). The abundance of each of these individual woody species provided better predictions of the hazel dormouse’s habitat selection than the combined summed abundance of hard mast species, soft mast species alone, or than overall summed abundance of woody species (Table S4).

**Figure 4.**
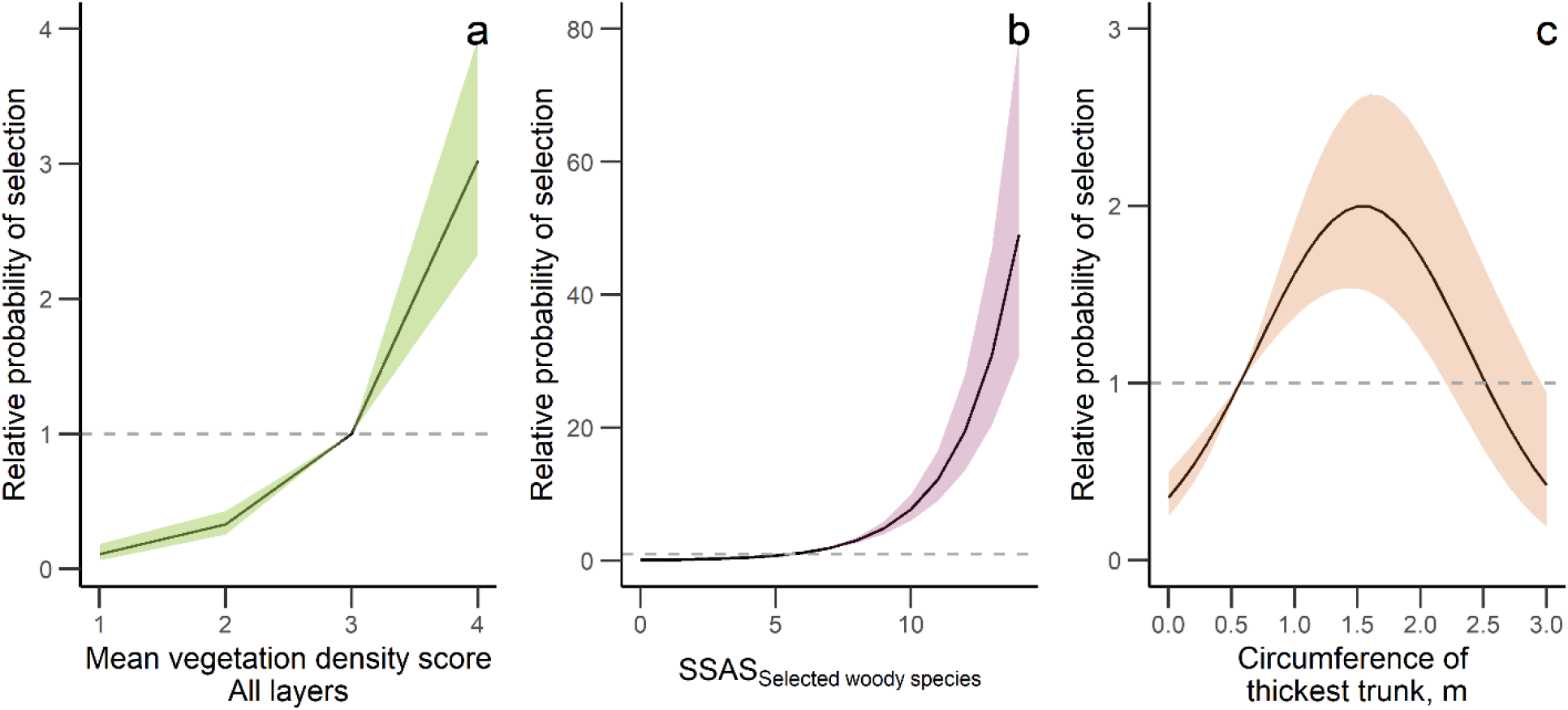
The predicted relationship ± 95% confidence interval between the relative probability of an area being selected by a hazel dormouse and (a) mean vegetation density score, (b) summed species abundance score (SSAS) of woody species selected by the dormice, and (c) circumference of the thickest trunk among 19 radio-tracked hazel dormice in a population located in a managed forest in Svanninge Bjerge, Denmark, 2013-2014. Horizontal lines indicate use = availability.

**Table 4.**
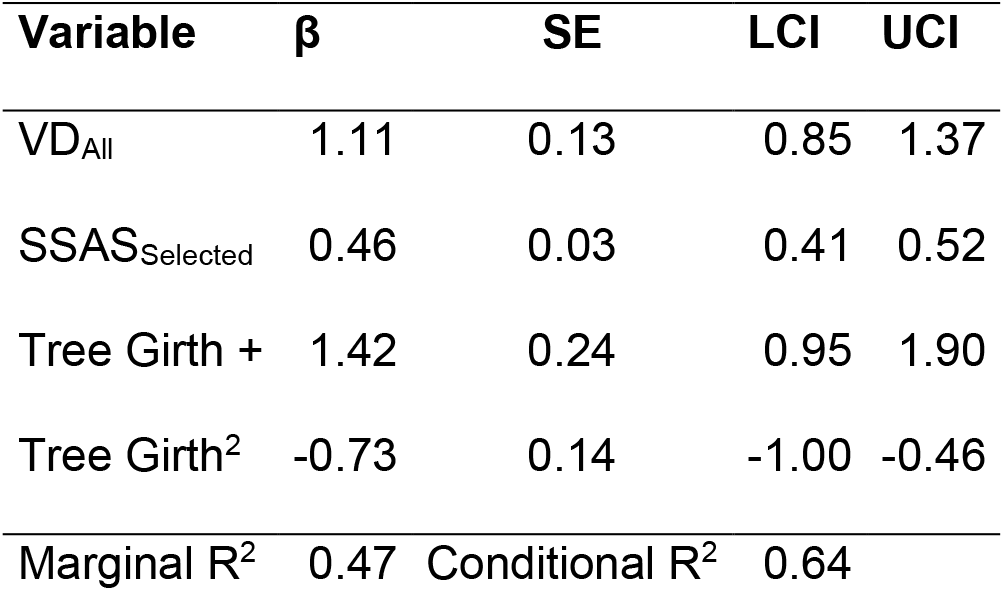
Selection coefficients (β), standard error (SE), lower (LCI) and upper (UCI) 95% confidence interval for the most parsimonious model (RSF) to explain nocturnal within home range habitat selection of 19 radio-tagged hazel dormice in Svanninge Bjerge, Denmark (n = 953 telemetry fixe and 1131 availability fixes).

| Variable | $\beta$ | SE | LCI | UCI |
| --- | --- | --- | --- | --- |
| VD<sub>All</sub> | 1.11 | 0.13 | 0.85 | 1.37 |
| SSAS<sub>Selected</sub> | 0.46 | 0.03 | 0.41 | 0.52 |
| Tree Girth + | 1.42 | 0.24 | 0.95 | 1.90 |
| Tree Girth<sup>2</sup> | -0.73 | 0.14 | -1.00 | -0.46 |
| Marginal R<sup>2</sup> | 0.47 | Conditional R<sup>2</sup> | 0.64 |  |

**Table 5.** Selection coefficients (β), standard error (SE), lower (LCI) and upper (UCI) 95% confidence interval for abundance scores (0-3) of individual woody plant species to explain within home range habitat selection of 19 radio-tagged hazel dormice in in Svanninge Bjerge, Denmark (n = 953 telemetry fixe and 1131 availability fixes).

| Variable | $\beta$ | SE | LCI | UCI |
| --- | --- | --- | --- | --- |
| Maple (*Acer* spp.) | 2.52 | 1.09 | 0.39 | 4.66 |
| Birch (*Betula* spp.) | 0.54 | 0.07 | 0.40 | 0.68 |
| Elder (*Sambucus* spp.) | 1.84 | 0.83 | 0.21 | 3.48 |
| Hazel (*Corylus avellana*) | 0.87 | 0.35 | 0.18 | 1.57 |
| Raspberry (*Rubus ideaus*) | 0.46 | 0.09 | 0.28 | 0.63 |
| Beech (*Fagus sylvatica*) | 0.53 | 0.06 | 0.41 | 0.65 |
| Willow (*Salix* spp.) | 0.26 | 0.12 | 0.02 | 0.50 |
| Blackberry (*Rubus plicatus*) | 0.25 | 0.07 | 0.11 | 0.38 |
| Rowan (*Sorbus* spp.) | 0.97 | 0.17 | 0.63 | 1.31 |
| Cherry (*Prunus* spp.) | 0.45 | 0.22 | 0.03 | 0.88 |
| Honeysuckle (*Lonicera* spp.) | 2.07 | 0.67 | 0.76 | 3.38 |
| Marginal R<sup>2</sup> | 0.50 | Conditional R<sup>2</sup> | 0.67 |  |

Habitat selection varied considerably among the tracked individuals. Selection for vegetation density (i.e. against openness) was stronger among individuals with more dense vegetation available within their home range and among larger individuals (Table 6, Fig. 5). Selection for the selected woody species was stronger among smaller individuals and individuals tracked earlier in the season (Table 6, Fig. 6). Selection for tree trunk circumference at breast-height did not vary among the tracked individuals (Table 6).

**Table 6.** Estimate (β), standard error (SE), lower (LCI) and upper (UCI) 95% confidence interval for explanatory variables to explain individual variation in selection for vegetation density, summed species abundance score (SSAS) of woody species selected by the dormice, and tree girth of 19 radio-tagged hazel dormice in a managed forest population located in Svanninge Bjerge, Denmark (n = 19 individuals).

| Variable | $\beta$ | SE | LCI | UCI |
| --- | --- | --- | --- | --- |
| *Selection for vegetation density* |  |  |  |  |
| Intercept | -11.78 | 3.16 | -17.98 | -5.58 |
| VD<sub>All</sub> | 2.07 | 0.82 | 0.46 | 3.67 |
| Mass | 0.35 | 0.06 | 0.24 | 0.46 |
| R<sup>2</sup> | 0.73 | R<sup>2</sup><sub>adjusted</sub> | 0.70 |  |
| *Selection for SSAS<sub>Selected</sub>* |  |  |  |  |
| Intercept | 7.34 | 1.49 | 4.43 | 10.26 |
| Mass | -0.05 | 0.02 | -0.07 | -0.02 |
| log(Julian day) | -1.16 | 0.27 | -1.69 | -0.63 |
| R<sup>2</sup> | 0.73 | R<sup>2</sup><sub>adjusted</sub> | 0.70 |  |
| *Selection for tree girth* |  |  |  |  |
| Intercept | 0.54 | 0.56 | -0.57 | 1.64 |
| R<sup>2</sup> | 0.00 | R<sup>2</sup><sub>adjusted</sub> | 0.00 |  |

**Figure 5.**
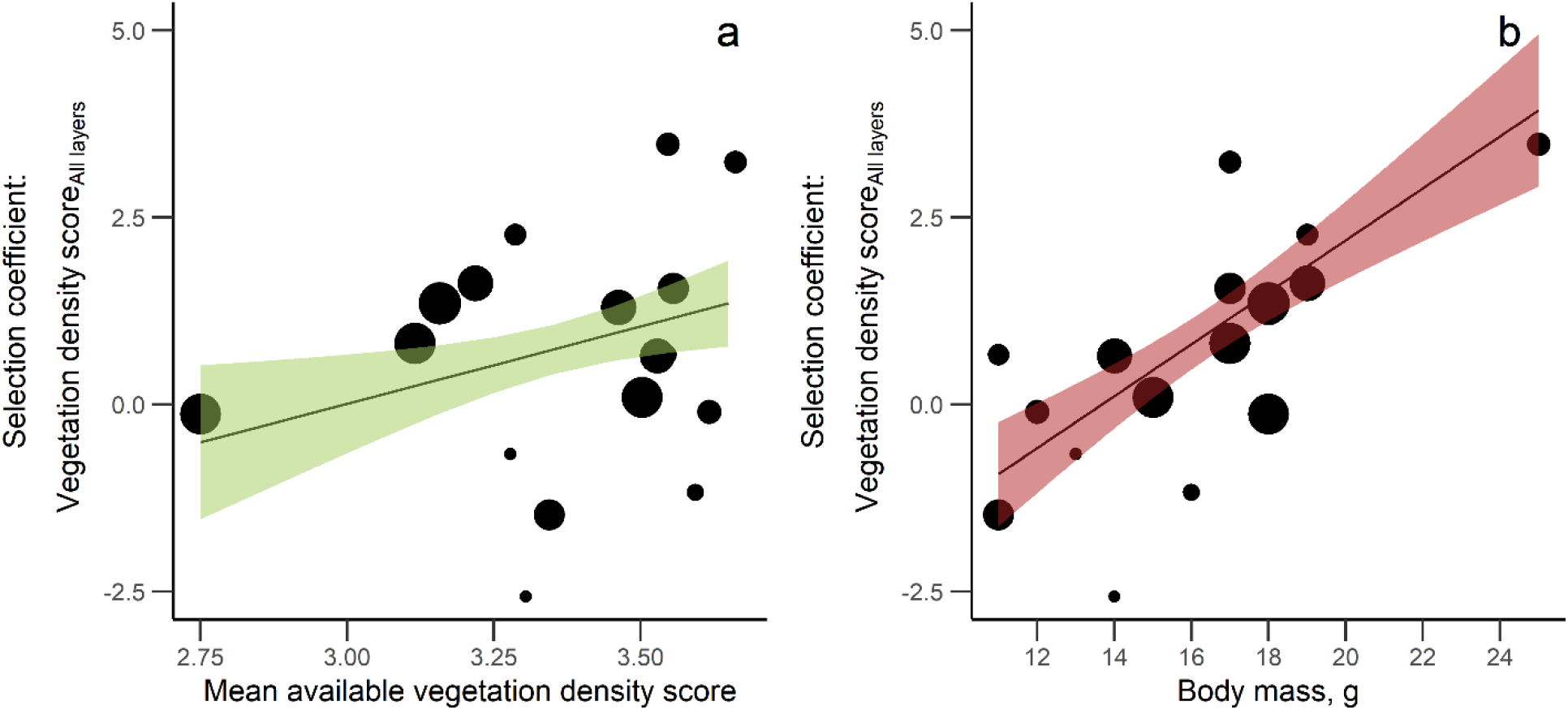
The predicted relationship ± 95% confidence interval between individual hazel dormice’s selection coefficients for vegetation density score and (a) mean vegetation density score within home range and (b) body mass among 19 radio-tracked hazel dormice in a population located in a managed forest in Svanninge Bjerge, Denmark, 2013-2014. Points represent selection coefficients of individuals. Point size indicate the inverse-variances which were used as weights in the analysis.

**Figure 6.**
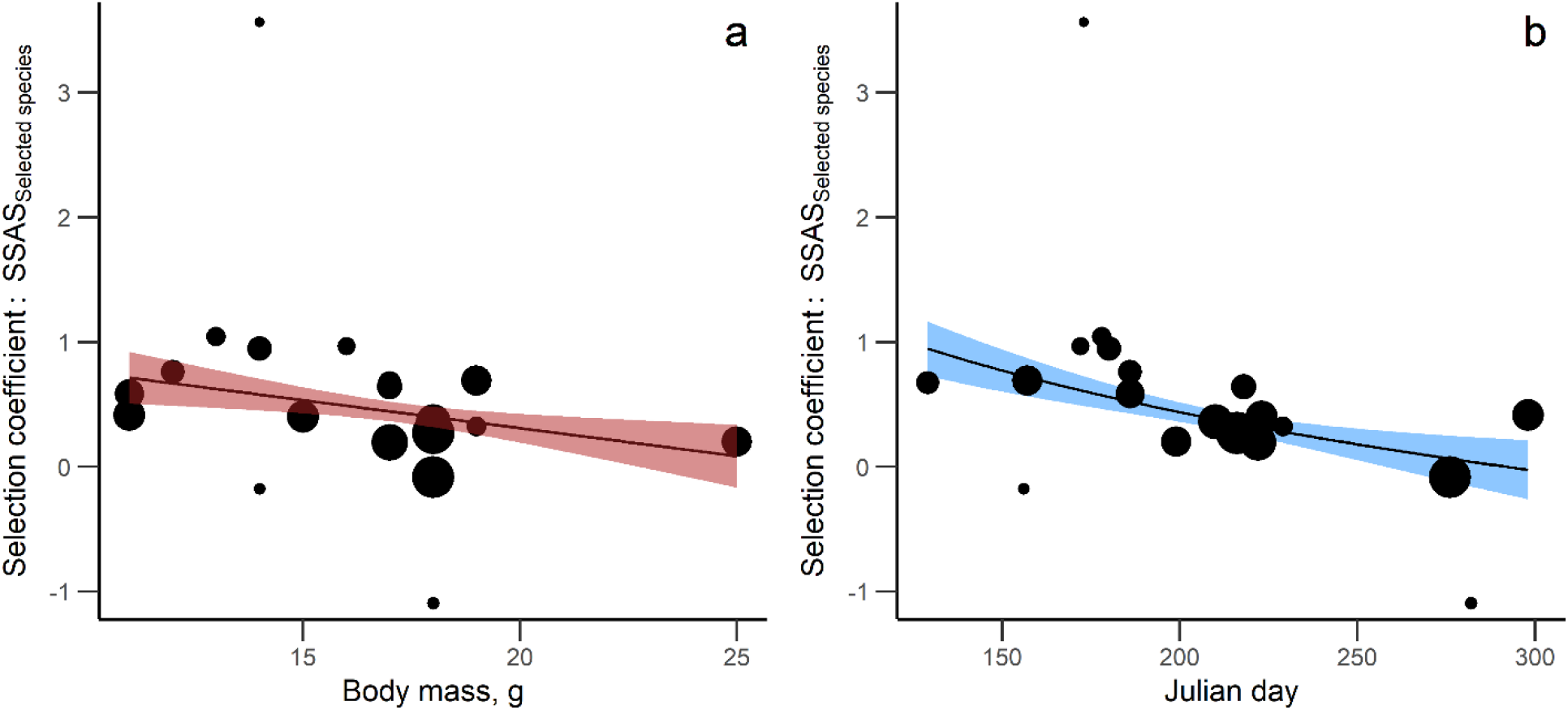
The predicted relationship ± 95% confidence interval between individual hazel dormice’s selection coefficients for summed species abundance score (SSAS) of woody species selected by the dormice and (a) body mass and (b) Julian day among 19 radio-tracked hazel dormice in a population located in a managed forest in Svanninge Bjerge, Denmark, 2013-2014. Points represent selection coefficients of individuals. Point size indicate the inverse-variances which were used as weights in the analysis.

## 4. Discussion

Our models provided unequivocal evidence with high predictive power that the hazel dormouse selected for specific woody plant species and dense and species rich tree vegetation on as well as within home range scale, whereas selection for forest vegetation age differed between the two spatial scales (the youngest tree stands most selected at home range level, stands with intermediate tree size most selected within home ranges). Our results highlight the importance of accessibility of a rich and diverse food base (provided by different species of woody plants) and high vegetation connectivity for movement and safety. Decreasing selection strengths with advancing calendar dates and increasing body mass further indicate that micro-habitat selection patterns correlate with energetic status and requirements. Our results emphasize targeted management of forest habitats as an important management action to ensure viable dormouse populations in a contiguous forest landscape. Most notably, this was illustrated by the predicted probability of occupancy between nest locations that varied from less than 1 to more than 95% as function of a single predictor variable (summed species abundance score of selected woody species). In reality, this means that forest managers with relatively simple means can improve habitat quality and carrying capacity for hazel dormice substantially by favouring species diverse, dense and not too mature tree stands within forests mainly managed for other purposes such as timber production.

Strong selection for summed abundance of a large number of woody plant species supports conclusions from other studies showing that a rich species composition is particularly important to satisfy the hazel dormouse’s requirements for resources and a continuous food supply of flowers, fruits, and invertebrates during its active season (Bright and Morris, 1996; Juškaitis and Baltrūnaitė, 2013; Juškaitis *et al*., 2016; Büchner *et al*., 2018; Goodwin *et al*., 2020). Although favoured vegetation types of the dormouse have shown to vary considerably between geographical locations (Juškaitis and Baltrūnaitė, 2013), stable food supplies throughout the active season is of paramount importance for hazel dormice (Bright and Morris, 1996). As the dormouse does not store food (Juškaitis, 2014a), it is critically dependent upon the timing of available food resources (Bright and Morris, 1996). Since different plant species have flowers and berries at different seasons, the strong positive correlation between occupancy and summed species abundance of all woody species makes perfect biological sense. Woody species that were selected at both spatial scales included beech, blackberry, elder, hazel, rowan, and willow which collectively may provide a good overlap in seasonal phenology of the various food objects throughout the dormouse’s active season. A good range of woody species may also be related to a high diversity and abundance of invertebrates which is believed to be critical for the dormouse, especially when the production of flowers, fruits, and nuts is scarce in the spring (Chanin *et al*., 2015). This supports other studies, indicating that the dormouse may not be such a selective feeder as thought in the past (Bright and Morris, 1996) but can occupy a wider variety of habitats (Trout *et al*., 2012; Juškaitis and Baltrūnaitė, 2013; Cartledge *et al*., 2021). This opportunistic adaptability to use food resources according to local species compositions may make it more robust to changes in species compositions caused by for example climate changes or changed forest management actions (Juškaitis *et al*., 2016; Goodwin *et al*., 2020). Within home ranges, we observed a higher selection strength for high abundance of selected woody species among smaller individuals and among individuals tracked earlier in the season, indicating the energetic constraints that affect the spatial decisions of these individuals as they have a higher demand for food resources to cover their energy expenditure after hibernation for growth and reproduction (Sozio *et al*., 2016). Other studies have found that the dormice’s use of torpor additionally may be a way to cope with varying food availability (Pretzlaff *et al*., 2014).

At both spatial scales, we observed a selection for vegetation density which enables safe movement options for the dormouse (Bright, 1998; Juškaitis *et al*., 2013). We found that high vegetation connectivity in the space below 10 m tree height was important for home range selection within populations, which resemble the dormouse’s preference for early successional woody habitats that naturally are species diverse and have a more complex vegetation structure (Swanson *et al*., 2011). The preference for dense vegetation might have been even higher if our study had included natural nests as Wolton (2009) found that where nesting conditions are good the hazel dormouse may prefer to build nests in unenclosed situations rather than in tree hollows and nest boxes. Hence, the presence of dormice may have been more unnoticed in very dense habitats of our study area. For habitat selection within home ranges, hazel dormice selected for vegetation density in all vertical layers which indicates the importance of high branch-connectivity both in terms of hiding from potential predators and for moving between different trees, shrubs, and other habitats (Bright, 1998; Juskaitis, 2004; Juškaitis *et al*., 2013; Dondina *et al*., 2016).

Selection strength for dense vegetation was stronger among individuals with higher average vegetation density available within their home ranges which shows how hazel dormice can adjust their movement behaviour according to the habitat composition within their home range. In home ranges with less amount of dense vegetation, because of for example forest fragmentation and intense forest practices, the dormice may not be able to fulfil their energetic requirements alone from the habitats they can encounter just by moving through dense vegetation but have to also perform crossings in more open vegetation. This shows that hazel dormice may be able to cope with minor habitat fragmentations, which other studies similarly report (Büchner, 2008; Mortelliti *et al*., 2013; Kelm *et al*., 2015), and suggest that habitat loss and poor habitat quality at landscape-level may be more critical for their conservation (Mortelliti *et al*., 2011; Goodwin *et al*., 2018a). The habitat quality of habitat patches has been shown to positively affect the survival and density of Italian hazel dormouse populations (Mortelliti *et al*., 2014), highlighting the importance of preserving areas with high habitat quality in the larger patch system, and support dispersal to these by improving connectivity of habitat patches through for example plantation of hedgerows (Dietz *et al*., 2018; Dondina *et al*., 2018). We also observed a higher selection strength for dense vegetation among larger individuals which implies the energetic constraints shaping their habitat selection (Gallagher *et al*., 2017; Mortensen *et al*., 2021). Smaller individuals may be more willing to expose themselves to risks and potential predators in order to increase their energy intake for reproduction, growth, and prepare for hibernation (Juškaitis *et al*., 2013; Pretzlaff *et al*., 2014).

We saw that home ranges within dormouse populations in managed woodlands were more often located in younger forest habitats, indicated by the selection for smaller tree circumference, whereas habitats with more intermediate tree circumferences were selected within home ranges of individuals. Both indicate the hazel dormouse’s requirements for dynamic mid-successional forest habitats (Goodwin *et al*., 2018a). Nest boxes have been found to attract dormice because of their resemblance to tree holes (Morris *et al*., 1990; Bright and Morris, 1991; Juškaitis, 2005), which may enhance the density of dormice in young forest habitats where natural tree holes are scarce (Vilhelmsen, 2003). However, as we show, dormice strongly select for woody pioneer species and their selection for early successional forest habitats may to a higher degree resemble this preference. On the other hand, in heavily managed woodlands where dense vegetation may be lacking because of frequent coppicing, clear-cutting, or over-grazing, nest boxes can be a management tool to improve conservation of hazel dormouse populations by providing safe resting and breeding places (Juškaitis, 2005). Our results show that we can increase the abundance of dormice considerably with targeted forest management practices. Disturbances from forest management practices may cause a decrease in population density in the short term, but the affected areas are typically recolonized within a short time (Trout *et al*., 2012; Sozio *et al*., 2016; Goodwin *et al*., 2018a; Juškaitis, 2020). Small-scale thinning and clear-cuts may even improve the quality of potential dormouse habitats by creating light open patches with structurally heterogeneous young shrubs (Berg, 1996; Wolton, 2009; Ramakers *et al*., 2014; Sozio *et al*., 2016; Juškaitis, 2020). Large-scale management practices can however be detrimental to the dormouse populations causing fragmentation, isolation, and loss of important forest patches of high quality (Mortelliti *et al*., 2011; Trout *et al*., 2012; Zapponi *et al*., 2013; Mortelliti *et al*., 2014; Sozio *et al*., 2016).

From a management point of view, our results indicate that the hazel dormouse even in the northern edge of its geographic distribution seems to occur at quite high densities under the right habitat conditions. With an average home range size of 0.5 ha in Denmark (R. M. Mortensen et al. unpubl. data) and social organisation with overlapping home ranges (Bright and Morris, 1991; Juškaitis *et al*., 2020), the habitats with the highest predicted occupancy (99%) must as minimum have sustained several individuals per hectare. This points directly to the targeted management of forest habitats as an potentially important management action to ensure viable dormouse populations in contiguous forest areas (Cartledge *et al*., 2021). The hazel dormouse’s dependence on a wide range of woody plants can be regarded as a management bonus as forest management aimed on improving living conditions for dormice can be combined with biodiversity considerations in general.

## 5. Conclusion

At home range level as well as within home ranges hazel dormice express a strong affinity for woody plant vegetation with high abundance-weighted species richness and high vegetation density. Selection for habitat parameters in general and variation in selection strengths as a function of date and body mass concede with existing knowledge on the species’ ecological requirements.

The incidence that the models with narrow confidence limits could predict more than 99% probability of home range occupancy under the most favourable combinations of habitat predictors demonstrates that the hazel dormouse has specific habitat requirements related to food and safety that should be possible to accommodate with relatively simple means in managed forests.

## Authors’ contributions

PS conceived the idea, raised funding, and managed the project. RMM, TBB, and PS developed the study design. RMM led the field work of the study with support from PS, MFF, and TBB. RMM performed the statistical analyses with support from PS. RMM and PS wrote the manuscript with support from MFF, LD, and TBB. All authors read and approved the final manuscript.

## Acknowledgements

This project originally constituted RMM’s MSc project at Aarhus University. We thank the landowners for letting us examine dormice presence and assess the habitats in their forests. Furthermore, we thank the Bikuben Foundation for letting us use their area for the radio telemetry. Thanks to Mads Kristian Warming, and the staff of Naturama for assisting with the radio telemetry, as well as to Peter Leth (the Danish Nature Agency) and Peter Mæhl (Ramboll) for letting us use their nest tubes from the national monitoring program in our analyses. Thanks to Ruth Morrison Svensson for reviewing the language of our paper. The project was financially supported by the Danish Nature Agency (Project title: “Forvaltning af hasselmus”).

## Data availability

The datasets used and analysed during the current study are available from doi.org/10.23642/usn.19425311 (will be activated upon acceptance of publication).

