## Supplemental material for "Hazel dormouse in managed woodland select for young, dense, and species-rich tree stands"

**S1:** Overview of radio tracked individuals in a hazel dormouse population in Svanninge Bjerge, Denmark, tracked in 2013-2014. No signal was obtained from dormouse #20 the evening after being tagged.

| ID | Sex | Weight, g | Tracking period | Tracking nights | Fixes |
|---|---|---|---|---|---|
| 1 | Male | 17 | 06/05/2013 - 13/05/2013 | 7 | 41 |
| 2 | Male | 19 | 05/06/2013 - 08/06/2013 | 3 | 33 |
| 3 | Male | 14 | 05/06/2013 - 05/06/2013 | 1 | 3 |
| 4 | Female | 25 | 16/07/2013 - 21/07/2013 | 6 | 82 |
| 5 | Male | 18 | 29/07/2013 - 31/07/2013 | 2 | 32 |
| 6 | Male | 18 | 02/08/2013 - 07/08/2013 | 6 | 90 |
| 7 | Male | 17 | 06/08/2013 - 07/08/2013 | 2 | 21 |
| 8 | Male | 15 | 09/08/2013 - 15/08/2013 | 6 | 55 |
| 9 | Male | 17 | 09/08/2013 - 13/08/2013 | 4 | 47 |
| 10 | Male | 19 | 16/08/2013 - 19/08/2013 | 3 | 43 |
| 11 | Female | 18 | 30/09/2013 - 08/10/2013 | 8 | 103 |
| 12 | Female | 18 | 09/10/2013 - 09/10/2013 | 1 | 3 |
| 13 | Female | 11 | 24/10/2013 - 27/10/2013 | 4 | 67 |
| 14 | Male | 14 | 21/06/2014 - 24/06/2014 | 3 | 36 |
| 15 | Female | 16 | 21/06/2014 - 22/06/2014 | 1 | 14 |
| 16 | Male | 13 | 25/06/2014 - 29/06/2014 | 4 | 64 |
| 17 | Female | 14 | 27/06/2014 - 03/07/2014 | 6 | 82 |
| 18 | Male | 12 | 03/07/2014 - 08/07/2014 | 5 | 70 |
| 19 | Female | 11 | 03/07/2014 - 08/07/2014 | 5 | 67 |
| 20 | Female | 13 | 10/06/2014 - 10/06/2014 | 0 | - |

**S2:** The model selection results for the best candidate models investigating hazel dormouse occupancy in nest boxes and nest tubes (n = 588) across hazel dormouse populations in fifteen managed forest patches in Denmark in 2013-2014. Models were ranked based on AICc. Most parsimonious model within ΔAICc < 2 in bold.

| Variables | df | log likelihood | AICc | $\Delta AIC_c$ | weight | $R^2_{marginal}$ | $R^2_{conditional}$ |
|---|---|---|---|---|---|---|---|
| 5+6+7+9 | 6 | -252.31 | 516.76 | 0.00 | 0.27 | 0.51 | 0.68 |
| **4+7+9** | **5** | **-253.42** | **516.94** | **0.18** | **0.24** | **0.49** | **0.65** |
| 5+6+7+8+9 | 7 | -252.20 | 518.60 | 1.84 | 0.11 | 0.50 | 0.67 |
| 4+7+8+9 | 6 | -253.36 | 518.86 | 2.10 | 0.09 | 0.49 | 0.65 |
| 1+2+7+9 | 6 | -253.40 | 518.95 | 2.19 | 0.09 | 0.49 | 0.65 |
| 1+2+3+7+9 | 7 | -253.22 | 520.63 | 3.87 | 0.04 | 0.49 | 0.65 |
| 1+2+7+8+9 | 7 | -253.35 | 520.89 | 4.13 | 0.03 | 0.49 | 0.65 |
| 4+7 | 4 | -256.65 | 521.36 | 4.60 | 0.03 | 0.49 | 0.64 |
| 4+7+8 | 5 | -255.95 | 522.00 | 5.24 | 0.02 | 0.49 | 0.64 |
| 1+2+3+7+8+9 | 8 | -253.21 | 522.66 | 5.91 | 0.01 | 0.49 | 0.64 |
| 5+6+7 | 5 | -256.50 | 523.10 | 6.34 | 0.01 | 0.50 | 0.67 |
| 5+6+7+8 | 6 | -255.48 | 523.11 | 6.35 | 0.01 | 0.50 | 0.66 |
| 1+2+7 | 5 | -256.59 | 523.29 | 6.53 | 0.01 | 0.49 | 0.65 |
| 1+2+7+8 | 6 | -255.94 | 524.02 | 7.26 | 0.01 | 0.49 | 0.64 |
| 2+7+9 | 5 | -257.19 | 524.49 | 7.73 | 0.01 | 0.46 | 0.64 |
| 2+3+7+9 | 6 | -256.23 | 524.60 | 7.84 | 0.01 | 0.46 | 0.63 |
| 1+2+3+7 | 6 | -256.35 | 524.85 | 8.09 | 0.00 | 0.49 | 0.64 |
| 1+7+9 | 5 | -257.51 | 525.12 | 8.36 | 0.00 | 0.46 | 0.61 |
| 1+2+3+7+8 | 7 | -255.87 | 525.92 | 9.16 | 0.00 | 0.49 | 0.64 |
| 2+7+8+9 | 6 | -257.17 | 526.48 | 9.72 | 0.00 | 0.46 | 0.63 |

*1 = $VD_{Low}$, 2 = $VD_{Middle}$, 3 = $VD_{High}$, 4 = $VD_{LM}$, 5 = $VD_{All}$ 6 = $VD_{All}^2$, 7 = $SSAS_{Selected}$, 8 = Height, 9 = Girth*

**S3:** The model selection results for the best candidate models investigating hazel dormouse occupancy in nest boxes and nest tubes (n = 588) across hazel dormouse populations in fifteen managed forest patches in Denmark in 2013-2014 as a function of woody species abundance within the assessment area. Most parsimonious model within ΔAICc < 2 in bold.

| Variables | df | log likelihood | AICc | $\Delta AIC_c$ | weight | $R^2_{marginal}$ | $R^2_{conditional}$ |
|---|---|---|---|---|---|---|---|
| 1+2+3+4+5+6+7+14+16+17+20 | 13 | -251.32 | 529.27 | 0.00 | 0.10 | 0.47 | 0.64 |
| 1+2+3+4+5+6+7+14+16+17+18+20 | 14 | -250.29 | 529.32 | 0.05 | 0.10 | 0.48 | 0.64 |
| 1+2+3+4+5+6+7+14+15+16+17+18+20 | 15 | -249.88 | 530.60 | 1.33 | 0.05 | 0.48 | 0.64 |
| 1+2+3+4+5+6+7+14+15+16+17+20 | 14 | -250.95 | 530.63 | 1.36 | 0.05 | 0.47 | 0.64 |
| 1+2+3+4+5+6+7+8+14+16+17+20 | 14 | -250.99 | 530.71 | 1.44 | 0.05 | 0.47 | 0.64 |
| 1+2+3+4+5+6+7+13+14+16+17+20 | 14 | -251.01 | 530.75 | 1.48 | 0.05 | 0.48 | 0.63 |
| **1+2+3+4+5+6+7+14+16+17** | **12** | **-253.11** | **530.76** | **1.49** | **0.05** | **0.47** | **0.64** |
| 1+2+3+4+5+6+7+11+14+16+17+20 | 14 | -251.03 | 530.79 | 1.52 | 0.05 | 0.47 | 0.64 |
| 1+2+3+4+5+6+7+13+14+16+17+18+20 | 15 | -250.03 | 530.90 | 1.63 | 0.05 | 0.48 | 0.64 |
| 1+2+3+4+5+6+7+11+14+16+17+18+20 | 15 | -250.05 | 530.94 | 1.67 | 0.05 | 0.48 | 0.64 |
| 1+2+3+4+5+6+7+14+16+17+18 | 13 | -252.17 | 530.98 | 1.71 | 0.04 | 0.47 | 0.64 |
| 1+2+3+4+5+6+7+8+14+16+17+18+20 | 15 | -250.08 | 531.01 | 1.74 | 0.04 | 0.48 | 0.64 |
| 1+2+3+4+5+6+7+12+14+16+17+18+20 | 15 | -250.11 | 531.05 | 1.78 | 0.04 | 0.48 | 0.63 |
| 1+2+3+4+5+6+7+12+14+16+17+20 | 14 | -251.18 | 531.10 | 1.83 | 0.04 | 0.48 | 0.63 |
| 1+2+3+4+5+6+7+9+14+16+17+20 | 14 | -251.23 | 531.20 | 1.92 | 0.04 | 0.47 | 0.64 |
| 1+2+3+4+5+6+7+10+14+16+17+20 | 14 | -251.25 | 531.24 | 1.97 | 0.04 | 0.47 | 0.64 |
| 1+2+3+4+5+6+7+9+14+16+17+18+20 | 15 | -250.20 | 531.24 | 1.97 | 0.04 | 0.48 | 0.64 |
| 1+2+3+4+5+6+7+14+16+17+19+20 | 14 | -251.27 | 531.28 | 2.01 | 0.04 | 0.47 | 0.64 |
| 1+2+3+4+5+6+7+14+16+17+18+19+20 | 15 | -250.24 | 531.32 | 2.05 | 0.04 | 0.48 | 0.64 |
| 1+2+3+4+5+6+7+10+14+16+17+18+20 | 15 | -250.25 | 531.34 | 2.07 | 0.04 | 0.48 | 0.64 |

*1 = Beech, 2 = Blackberry, 3 = Elder, 4 = Hawthorne, 5 = Larch, 6 = Rowan, 7 = Willow, 8 = Maple, 9 = Ash, 10 = Birch, 11 = Oak, 12 = Alder, 13 = Elm, 14 = Pine, 15 = Honeysuckle, 16 = Spruce, 17 = Hazel, 18 = Raspberry, 19 = Cherry, 20 = Thuja*

**S4:** The model selection results for the best candidate models investigating habitat selection within home ranges of nineteen hazel dormice (n = 2084) in a managed forest population located in Svanninge Bjerge, Denmark. Most parsimonious model within ΔAICc < 2 in bold.

| Variables | df | log likelihood | AICc | $\Delta AIC_c$ | weight | $R^2_{marginal}$ | $R^2_{conditional}$ |
|---|---|---|---|---|---|---|---|
| **6+8+11+12** | **5** | **-876.29** | **1762.61** | **0.00** | **0.60** | **0.41** | **0.65** |
| 1+2+3+8+11+12 | 7 | -874.88 | 1763.82 | 1.21 | 0.33 | 0.40 | 0.64 |
| 1+3+8+11+12 | 6 | -877.45 | 1766.94 | 4.33 | 0.07 | 0.35 | 0.63 |
| 5+8+11+12 | 5 | -882.13 | 1774.30 | 11.69 | 0.00 | 0.39 | 0.65 |
| 2+3+8+11+12 | 6 | -881.87 | 1775.78 | 13.17 | 0.00 | 0.39 | 0.62 |
| 3+8+11+12 | 5 | -887.40 | 1784.83 | 22.22 | 0.00 | 0.37 | 0.65 |
| 6+8+10 | 4 | -889.35 | 1786.72 | 24.11 | 0.00 | 0.39 | 0.62 |
| 6+8 | 3 | -893.93 | 1793.86 | 31.25 | 0.00 | 0.38 | 0.65 |
| 1+2+3+8 | 5 | -892.68 | 1795.38 | 32.77 | 0.00 | 0.38 | 0.62 |
| 5+8 | 3 | -900.62 | 1807.24 | 44.63 | 0.00 | 0.38 | 0.64 |
| 4+8+11+12 | 5 | -898.99 | 1808.01 | 45.40 | 0.00 | 0.35 | 0.63 |
| 1+3+8 | 4 | -900.24 | 1808.50 | 45.89 | 0.00 | 0.35 | 0.63 |
| 2+3+8 | 4 | -900.53 | 1809.07 | 46.46 | 0.00 | 0.36 | 0.59 |
| 1+2+8+11+12 | 6 | -898.95 | 1809.94 | 47.32 | 0.00 | 0.34 | 0.62 |
| 2+8+11+12 | 5 | -903.53 | 1817.10 | 54.49 | 0.00 | 0.34 | 0.60 |
| 1+8+11+12 | 5 | -904.46 | 1818.95 | 56.34 | 0.00 | 0.37 | 0.62 |
| 8+9+11+12 | 5 | -904.70 | 1819.44 | 56.83 | 0.00 | 0.36 | 0.62 |
| 6+7+11+12 | 5 | -905.46 | 1820.95 | 58.34 | 0.00 | 0.37 | 0.61 |
| 1+2+3+7+11+12 | 7 | -905.27 | 1824.60 | 61.99 | 0.00 | 0.40 | 0.61 |
| 8+11+12 | 4 | -912.39 | 1832.80 | 70.19 | 0.00 | 0.37 | 0.63 |

*1 = $VD_{Low}$, 2 = $VD_{Middle}$, 3 = $VD_{High}$, 4 = $VD_{LM}$, 5 = $VD_{MH}$, 6 = $VD_{All}$, 7 = $SSAS_{Softmast}$, 8 = $SSAS_{Selected}$, 9 = Tree canopy cover, 10 = Height, 11 = Girth, 12 = $Girth^2$*

**S5:** The model selection results for the best candidate models investigating habitat selection within home ranges of nineteen hazel dormice (n = 2084) in a managed forest population located in Svanninge Bjerge, Denmark as a function of woody species abundance within the assessment area. Most parsimonious model within ΔAICc < 2 in bold.

| Variables | df | log likelihood | AICc | $\Delta AIC_c$ | weight | $R^2_{marginal}$ | $R^2_{conditional}$ |
|---|---|---|---|---|---|---|---|
| 1+2+3+4+5+6+7+8+9+10+12+14 | 13 | -937.73 | 1901.63 | 0.00 | 0.19 | 0.50 | 0.66 |
| **1+2+3+5+6+7+8+9+10+12+14** | **12** | **-939.04** | **1902.23** | **0.60** | **0.14** | **0.50** | **0.67** |
| 1+2+3+4+5+6+7+8+9+10+11+12+14 | 14 | -937.48 | 1903.16 | 1.53 | 0.09 | 0.50 | 0.67 |
| 1+2+3+4+5+6+7+8+9+10+12+13+14 | 14 | -937.65 | 1903.51 | 1.88 | 0.07 | 0.50 | 0.66 |
| 1+2+3+5+6+7+8+9+10+11+12+14 | 13 | -938.69 | 1903.56 | 1.93 | 0.07 | 0.50 | 0.67 |
| 1+2+3+4+5+6+7+8+9+10+14 | 12 | -939.97 | 1904.10 | 2.47 | 0.06 | 0.50 | 0.66 |
| 1+2+3+4+5+6+7+8+10+12+14 | 12 | -939.98 | 1904.11 | 2.48 | 0.06 | 0.50 | 0.67 |
| 1+2+3+5+6+7+8+9+10+12+13+14 | 13 | -938.98 | 1904.13 | 2.50 | 0.05 | 0.50 | 0.67 |
| 1+2+3+5+6+7+8+9+10+14 | 11 | -941.31 | 1904.74 | 3.11 | 0.04 | 0.50 | 0.67 |
| 1+2+3+5+6+7+8+10+12+14 | 11 | -941.54 | 1905.22 | 3.58 | 0.03 | 0.50 | 0.67 |
| 1+2+3+5+6+7+8+9+10+11+12+13+14 | 14 | -938.65 | 1905.51 | 3.87 | 0.03 | 0.50 | 0.67 |
| 1+2+3+4+5+6+7+8+10+11+12+14 | 13 | -939.71 | 1905.60 | 3.97 | 0.03 | 0.50 | 0.67 |
| 1+2+3+4+5+6+7+8+9+10+11+14 | 13 | -939.74 | 1905.66 | 4.03 | 0.03 | 0.50 | 0.66 |
| 1+2+3+4+5+6+7+8+10+12+13+14 | 13 | -939.92 | 1906.01 | 4.38 | 0.02 | 0.50 | 0.67 |
| 1+2+3+4+5+6+7+8+9+10+13+14 | 13 | -939.92 | 1906.02 | 4.39 | 0.02 | 0.50 | 0.66 |
| 1+2+3+5+6+7+8+9+10+11+14 | 12 | -940.98 | 1906.10 | 4.47 | 0.02 | 0.50 | 0.67 |
| 1+2+3+5+6+7+8+10+11+12+14 | 12 | -941.15 | 1906.45 | 4.82 | 0.02 | 0.50 | 0.67 |
| 1+2+3+5+6+7+8+9+10+13+14 | 12 | -941.27 | 1906.68 | 5.05 | 0.02 | 0.50 | 0.67 |
| 1+2+3+5+6+7+8+10+12+13+14 | 12 | -941.49 | 1907.13 | 5.50 | 0.01 | 0.50 | 0.67 |
| 1+2+3+4+5+6+7+8+10+11+12+13+14 | 14 | -939.66 | 1907.53 | 5.90 | 0.01 | 0.50 | 0.67 |

*1 = Beech, 2 = Birch, 3 = Blackberry, 4 = Buckthorn, 5 = Elder, 6 = Hazel, 7 = Maple, 8 = Raspberry, 9 = Willow, 10 = Rowan, 11 = Oak, 12 = Cherry, 13 = Hawthorn, 14 = Honeysuckle*

**S6:** The model selection results for the best candidate models investigating nineteen dormice's individual selection coefficients for mean vegetation density in a managed forest population located in Svanninge Bjerge, Denmark in 2013-2014. Most parsimonious model within ΔAICc < 2 in bold.

| **Variables** | **df** | **log likelihood** | **AICc** | **$\Delta AIC_c$** | **weight** | **$R^2$** | **$R^2_{adjusted}$** |
|---|---|---|---|---|---|---|---|
| 1+2+6 | 5 | -30.94 | 76.49 | 0.00 | 0.17 | 0.78 | 0.74 |
| **1+6** | **4** | **-32.85** | **76.55** | **0.07** | **0.16** | **0.73** | **0.70** |
| 1+6+7 | 5 | -30.99 | 76.59 | 0.10 | 0.16 | 0.78 | 0.74 |
| 1+2+6+7 | 6 | -29.15 | 77.30 | 0.81 | 0.11 | 0.82 | 0.77 |
| 1+3+6+7 | 6 | -29.95 | 78.90 | 2.42 | 0.05 | 0.80 | 0.75 |
| 1+3+6 | 5 | -32.28 | 79.17 | 2.68 | 0.04 | 0.75 | 0.70 |
| 2+3+6 | 5 | -32.43 | 79.48 | 2.99 | 0.04 | 0.74 | 0.69 |
| 1+5+6 | 5 | -32.50 | 79.62 | 3.14 | 0.03 | 0.74 | 0.69 |
| 1+4+6+7 | 6 | -30.38 | 79.76 | 3.27 | 0.03 | 0.79 | 0.73 |
| 1+4+6 | 5 | -32.73 | 80.07 | 3.59 | 0.03 | 0.74 | 0.68 |
| 1+2+4+6 | 6 | -30.60 | 80.21 | 3.72 | 0.03 | 0.79 | 0.73 |
| 2+6 | 4 | -34.76 | 80.38 | 3.89 | 0.02 | 0.67 | 0.63 |
| 1+2+3+6 | 6 | -30.69 | 80.38 | 3.90 | 0.02 | 0.79 | 0.73 |
| 1+2+5+6 | 6 | -30.90 | 80.80 | 4.32 | 0.02 | 0.78 | 0.72 |
| 2+4+6 | 5 | -33.10 | 80.81 | 4.33 | 0.02 | 0.73 | 0.67 |
| 1+5+6+7 | 6 | -30.96 | 80.91 | 4.43 | 0.02 | 0.78 | 0.72 |
| 2+6+7 | 5 | -33.37 | 81.35 | 4.87 | 0.01 | 0.72 | 0.66 |
| 2+3+6+7 | 6 | -31.67 | 82.34 | 5.85 | 0.01 | 0.76 | 0.70 |
| 1+2+5+6+7 | 7 | -29.08 | 82.35 | 5.87 | 0.01 | 0.82 | 0.75 |
| 1+2+3+6+7 | 7 | -29.14 | 82.47 | 5.98 | 0.01 | 0.82 | 0.75 |

*1 = meanVD$_{All}$, 2 = meanSSAS$_{Selected}$, 3 = meanGirth, 4 = meanHeight, 5 = Sex, 6 = Mass, 7 = log(Julian day)*

**S7:** The model selection results for the best candidate models investigating nineteen dormice's individual selection coefficients for summed species abundance score of woody species selected by the dormice in a managed forest population located in Svanninge Bjerge, Denmark in 2013-2014. Most parsimonious model within ΔAICc < 2 in bold.

| Variables | df | log likelihood | AICc | $\Delta AIC_c$ | weight | $R^2$ | $R^2_{adjusted}$ |
|---|---|---|---|---|---|---|---|
| **6+7** | **4** | **1.39** | **8.08** | **0.00** | **0.41** | **0.73** | **0.70** |
| 4+6+7 | 5 | 2.13 | 10.36 | 2.28 | 0.13 | 0.75 | 0.70 |
| 3+6+7 | 5 | 1.98 | 10.65 | 2.57 | 0.11 | 0.75 | 0.70 |
| 2+6+7 | 5 | 1.46 | 11.70 | 3.62 | 0.07 | 0.74 | 0.68 |
| 5+6+7 | 5 | 1.40 | 11.82 | 3.74 | 0.06 | 0.73 | 0.68 |
| 1+6+7 | 5 | 1.40 | 11.82 | 3.74 | 0.06 | 0.73 | 0.68 |
| 2+4+6+7 | 6 | 2.87 | 13.25 | 5.17 | 0.03 | 0.77 | 0.71 |
| 2+3+6+7 | 6 | 2.68 | 13.64 | 5.56 | 0.03 | 0.77 | 0.70 |
| 1+4+6+7 | 6 | 2.16 | 14.69 | 6.61 | 0.01 | 0.75 | 0.68 |
| 4+5+6+7 | 6 | 2.13 | 14.74 | 6.66 | 0.01 | 0.75 | 0.68 |
| 3+5+6+7 | 6 | 2.10 | 14.79 | 6.71 | 0.01 | 0.75 | 0.68 |
| 1+3+6+7 | 6 | 2.05 | 14.89 | 6.81 | 0.01 | 0.75 | 0.68 |
| 1+2+6+7 | 6 | 1.51 | 15.97 | 7.89 | 0.01 | 0.74 | 0.66 |
| 2+5+6+7 | 6 | 1.49 | 16.02 | 7.94 | 0.01 | 0.74 | 0.66 |
| 1+5+6+7 | 6 | 1.40 | 16.20 | 8.12 | 0.01 | 0.73 | 0.66 |
| 1+2+3+6+7 | 7 | 3.91 | 16.36 | 8.28 | 0.01 | 0.80 | 0.72 |
| 1+2+4+6+7 | 7 | 3.52 | 17.14 | 9.06 | 0.00 | 0.79 | 0.71 |
| 2+3+5+6+7 | 7 | 3.51 | 17.17 | 9.09 | 0.00 | 0.79 | 0.71 |
| 7 | 3 | -5.12 | 17.83 | 9.75 | 0.00 | 0.47 | 0.44 |
| 2+4+5+6+7 | 7 | 2.98 | 18.23 | 10.15 | 0.00 | 0.78 | 0.69 |

*1 = meanVD$_{All}$, 2 = meanSSAS$_{Selected}$, 3 = meanGirth, 4 = meanHeight, 5 = Sex, 6 = Mass, 7 = log(Julian day)*

**S8:** The model selection results for the best candidate models investigating nineteen dormice's individual selection coefficients for circumference of thickest trunk in a managed forest population located in Svanninge Bjerge, Denmark in 2013-2014. Most parsimonious model within ΔAICc < 2 in bold.

| Variables | df | log likelihood | AICc | $\Delta AIC_c$ | weight | $R^2$ | $R^2_{adjusted}$ |
|---|---|---|---|---|---|---|---|
| 6 | -67.06 | 141.72 | 0.00 | 0.17 | 0.18 | 0.14 | -67.06 |
| **0** | **-68.99** | **142.74** | **1.01** | **0.10** | **0.00** | **0.00** | **-68.99** |
| 3+6 | -66.17 | 143.19 | 1.47 | 0.08 | 0.26 | 0.16 | -66.17 |
| 3 | -67.87 | 143.35 | 1.62 | 0.08 | 0.11 | 0.06 | -67.87 |
| 2 | -67.91 | 143.41 | 1.69 | 0.07 | 0.11 | 0.06 | -67.91 |
| 5+6 | -66.36 | 143.59 | 1.86 | 0.07 | 0.24 | 0.15 | -66.36 |
| 1+6 | -66.42 | 143.69 | 1.97 | 0.06 | 0.24 | 0.14 | -66.42 |
| 2+6 | -66.65 | 144.17 | 2.44 | 0.05 | 0.22 | 0.12 | -66.65 |
| 6+7 | -66.68 | 144.22 | 2.50 | 0.05 | 0.22 | 0.12 | -66.68 |
| 4+6 | -66.91 | 144.68 | 2.96 | 0.04 | 0.20 | 0.10 | -66.91 |
| 7 | -68.77 | 145.14 | 3.42 | 0.03 | 0.02 | -0.03 | -68.77 |
| 4 | -68.90 | 145.40 | 3.68 | 0.03 | 0.01 | -0.05 | -68.90 |
| 5 | -68.93 | 145.45 | 3.73 | 0.03 | 0.01 | -0.05 | -68.93 |
| 1 | -68.99 | 145.59 | 3.86 | 0.03 | 0.00 | -0.06 | -68.99 |
| 1+3 | -67.58 | 146.01 | 4.29 | 0.02 | 0.14 | 0.03 | -67.58 |
| 2+3 | -67.64 | 146.13 | 4.41 | 0.02 | 0.13 | 0.02 | -67.64 |
| 3+7 | -67.70 | 146.25 | 4.53 | 0.02 | 0.13 | 0.02 | -67.70 |
| 3+6+7 | -65.85 | 146.32 | 4.60 | 0.02 | 0.28 | 0.14 | -65.85 |
| 3+5 | -67.78 | 146.42 | 4.69 | 0.02 | 0.12 | 0.01 | -67.78 |
| 1+2+6 | -65.97 | 146.55 | 4.83 | 0.02 | 0.27 | 0.13 | -65.97 |

*0 = null model, 1 = meanVD$_{All}$, 2 = meanSSAS$_{Selected}$, 3 = meanGirth, 4 = meanHeight, 5 = Sex, 6 = Mass, 7 = log(Julian day)*
